# Effect of senolytic drugs in young female mice chemically induced to estropause

**DOI:** 10.1101/2024.05.22.595355

**Authors:** Bianca M Ávila, Bianka M Zanini, Karina P Luduvico, Thais L Oliveira, Jéssica D Hense, Driele N Garcia, Juliane Prosczek, Francielle M Stefanello, Pedro H da Cruz, Janice L Giongo, Rodrigo A Vaucher, Jeffrey B Mason, Michal M Masternak, Augusto Schneider

## Abstract

**Aims:** This study aimed to assess metabolic responses and senescent cell burden in young female mice induced to estropause and treated with senolytic drugs.

**Main methods:** Estropause was induced by 4-vinylcyclohexene diepoxide (VCD) injection in two-month-old mice. The senolytics dasatinib and quercetin (D+Q) or fisetin were given by oral gavage once a month from five to 11 months of age.

**Key findings:** VCD-induced estropause led to increased body mass and reduced albumin concentrations compared to untreated cyclic mice, without affecting insulin sensitivity, lipid profile, liver enzymes, or total proteins. Estropause decreased catalase activity in adipose tissue but had no significant effect on other redox parameters in adipose and hepatic tissues. Fisetin treatment reduced ROS levels in the hepatic tissue of estropause mice. Estropause did not influence senescence-associated beta-galactosidase activity in adipose and hepatic tissues, but increased senescent cell markers and fibrosis in ovaries. Senolytic treatment did not decrease ovarian cellular senescence induced by estropause.

**Significance:** Overall, the findings suggest that estropause leads to minor metabolic changes in young females, and senolytics had no protective effects despite increased ovarian senescence.

## INTRODUCTION

Reproductive aging entails the gradual and continuous decline of ovarian reserve until its exhaustion and the onset of menopause (Broekmans et al., 2009; Finch, 2014; Cavalcante et al., 2023). Menopause results in decreased estrogen levels, leading to metabolic changes such as increased body weight gain, adiposity and dyslipidemia, increasing the risk of chronic diseases (Selbac et al., 2018; Bitto et al., 2009; Lizcano and Guzmán, 2014). Several studies propose that menopause induces a systemic pro-oxidant state due diminished antioxidant capacity (Doshi et al., 2013; Bourgonje et al., 2020; Secomandi et al., 2022), which may contribute to metabolic changes observed. In mice, cessation of cyclicity, or estropause, can be chemically induced by 4-vinylcyclohexene diepoxide (VCD), an ovotoxic drug that causes ovarian failure, similar to the gradual process of follicle loss observed in peri-menopausal women (Chen et al., 2014; Romero-Aleshire et al., 2009).

Cellular senescence is a response to endogenous and exogenous stressors during aging, aiming to inhibit proliferation of aged or damaged cells, resulting in a state of permanent cell cycle arrest (Calcinotto et al., 2019; Secomandi et al., 2022). Cellular senescence plays important physiological roles in tissue homeostasis and tumor suppression (Hanahan and Weinberg, 2011; McHugh and Gil, 2018; Secomandi et al., 2022; Kudlova et al., 2022). However, the accumulation of senescent cells during aging can generate chronic low-grade inflammation (Song et al., 2020), as senescent cells secrete a variety of pro-inflammatory chemokines, cytokines, and interleukins, known as the senescence-associated secretory phenotype (SASP). This accumulation of senescent cells and the associated pro-inflammatory profile can trigger tissue dysfunction and age-related chronic diseases (Secomandi et al., 2022; Kudlova et al., 2022). Although age-related senescent cells burden is well-established, little is known about the role of senescence in the pathogenesis of diseases in females after menopause/estropause independent of age.

In this context, drugs with senolytic potential have been used to promote selective clearance of senescent cells (Kudlova et al., 2022; Yousefzadeh et al., 2018; Xu et al., 2018). Among these, fisetin and a combination of dasatinib and quercetin (D+Q) were shown to delay age-associated disorders and increase lifespan of older mice (Baker et al., 2011; Yousefzadeh et al., 2018; Xu et al., 2018). Despite this, the effects of senolytic drugs in young female mice seems minor and even negative (Fang et al., 2023; Garcia et al., 2024). However, D+Q treatment can reduce senescent cell markers in young mice ovaries (Hense et al., 2022; Garcia et al., 2024). Nevertheless, it is necessary to understand whether estropause increases senescent cell burden independent of age and if senolytic drugs have a protective role. Therefore, we aimed to evaluate the response of young estropause-induced females to senolytic drugs.

## METHODS

### Animals and treatments

This study was approved by the Universidade Federal de Pelotas Animal Experimentation Ethics Committee (018232/2021-13). All animal experimentation followed the National Council for the Control of Animal Experimentation (Brazil) and the ARRIVE guidelines. Female C57BL/6 mice were kept under controlled conditions of temperature (22 ± 2 °C), light (12-hour light/12-hour dark cycles) and humidity (40%-60%). The chemical estropause induction protocol was initiated in 60-day old females (n=33) receiving i.p. injections of VCD (160 mg/kg) diluted in sesame oil for 20 consecutive days (Brooks et al., 2016; Lohff et al., 2005; Romero-Aleshire et al., 2009; Ávila et al., 2023). Control females (n=27) received i.p. injections of vehicle during the same period.

Estropause was confirmed by vaginal cytology at 5 months of age and the treatment with senolytics was started two weeks later. Estropausal and cyclic female mice were divided into three groups each: placebo, fisetin and D+Q. The treatment was performed via gavage once a month for 3 consecutive days, for 6 months. The doses were 100mg/kg of body mass for fisetin (Yousefzadeh et al., 2018) and 5mg/kg of dasatinib and 50 mg/kg of quercetin (Xu et al., 2018). Both treatments were diluted in the vehicle composed of ethanol, PEG 400 and phosal. Mice in the placebo groups received 100 µL of vehicle via gavage.

In the week before euthanasia mice were subjected to an insulin tolerance test (ITT). At the end of the experiment mice were fasted for 4 hours, subjected to anesthesia with isoflurane and euthanized by exsanguination. Mice were dissected to remove ovarian, visceral adipose and liver tissues, which were flash frozen in nitrogen and stored in at −80° C. Ovarian tissue was also preserved in formaldehyde for histological analysis. Visceral body fat was dissected and weighed on a precision scale immediately after euthanasia.

### Vaginal cytology

Vaginal cytology was performed daily for one week to observe if females were already acyclic, according to the previous protocol (McLean et al., 2012). For the vaginal cell sample collection, a micropipette was positioned at the opening of the vaginal canal to perform a lavage. The vaginal lavage was placed on a slide and stained using rapid panoptic. The slides were observed under an optical microscope to verify estrous cycle stage of each mice (McLean et al., 2012).

### Insulin Tolerance Test (ITT)

The ITT was performed by injecting insulin (1 IU/kg of body mass) i.p. after 2 hours fasting early in the morning. Blood was collected through a small incision at the tip of the tail to check blood glucose levels with a glucometer (AccuChek Activ, Roche Diagnostics®, USA) at the following times: 0, 15, 30 and 60 minutes after injection (Fang et al., 2013).

### Ovarian follicle count

The ovaries were removed from 10% buffered formalin, dehydrated in alcohol, cleared in xylene and embedded in paraffin and were sequentially cut at a thickness of 5μm on a semi-automatic microtome (RM2245 38, Leica Biosystems Newcastle Ltd, Newcastle Upon Tyne, UK).

One in every six sections were selected and placed on slides. After drying in an oven at 56°C for 24 hours, the slides were stained with hematoxylin-eosin and mounted with coverslips and synthetic resin (Sigma®, St. Louis, MO, USA). The images of the ovarian sections were captured by a digital camera (Moticam 5.0, Motic®, Hong Kong, China) coupled to a microscope (Nikon Eclipse E200, Nikon Corporation, Japan), using 4, 10 and 40X objectives. Follicles with clearly visible oocyte nuclei were quantified. The final number of follicles was divided by the total number of sections from each ovary (Myers et al., 2004).

### Biochemical Analysis

Blood samples were collected at euthanasia allowed to clot for 20 min at room temperature and centrifuged at 1500 x g to separate serum. The concentrations of total cholesterol (TC), high-density cholesterol (HDL), triglycerides (TRI), albumin (ALB), total proteins (PT), alanine aminotransferase (ALT) and aspartate aminotransferase (AST) were evaluated in serum using commercial kits (Bioclin, Belo Horizonte, Minas Gerais and Labtest Virtue Diagnostic, Lagoa Santa, Minas Gerais) in an automated biochemical analysis equipment (Cobas Mira S, Roche®, Basel, Switzerland).

### Tissue protein extraction and quantification

Liver tissue was homogenized (1/10 w/v) using 20 mM sodium phosphate buffer (pH 7.4) containing 140 mM KCl. Adipose tissue was homogenized (1/10 w/v) using RIPPA buffer (pH 7.5) containing 50 mM Tris HCl, 150 mM NaCl and 150 mM EDTA. The homogenates were centrifuged at 3500 x g at 4°C for 10 minutes and the supernatants used in oxidative stress analysis. Protein was measured using the BCA Protein Assay kit (Thermo Scientific™), according to the manufacturer’s instructions.

### Reactive oxygen species (ROS) levels

ROS levels were determined by oxidation of DCFH-DA to fluorescent 2’,7’-dichlorofluorescein (DCF). DCF fluorescence intensity was recorded at 525 and 488 nm excitation 30 min after addition of DCFH-DA to the medium. ROS formation was expressed as µmol DCF/mg of protein (Ali et al., 1992).

### Total sulfhydryl content

The total thiol content was determined by the reduction of DTNB by thiols, forming a derivative compound (TNB), which absorption is measured in the spectrophotometer at 412 nm. The results were expressed as nmol TNB/mg of protein (Aksenov and Markesbery, 2001).

### Substances reactive to thiobarbituric acid (TBARS) levels

Lipid peroxidation was determined by estimating malondialdehyde (MDA) formation. TBA and trichloroacetic acid were added to the supernatant for the reaction. TBARS levels were quantified with absorbance at 535 nm and reported as nmol TBARS/mg of protein (Esterbauer and Cheeseman, 1990).

### Nitrites levels

The nitrite reaction was performed with sulfanilamide and N-1-naphthylethylenediamine dihydrochloride (NED), using the Griess reaction. Absorbance was measured at 540 nm. The results were expressed as μM nitrite/mg of protein (Stuehr and Nathan, 1989).

### Catalase (CAT) activity

CAT activity was measured based on the decomposition of H_2_O_2_ at 240 nm. CAT activity was reported as units/mg protein (Aebi, 1984).

### Superoxide dismutase (SOD) activity

SOD activity was measured based on the inhibition of superoxide-dependent adrenaline auto-oxidation by SOD in the sample at 480 nm. SOD activity was expressed in units/mg of protein (Misra and Fridovich, 1972).

### Senescence-associated beta-galactosidase (SA-βGAL) activity

The analysis of SA-βGAL activity was performed in hepatic and adipose tissue through spectrophotometry using the mammalian beta-galactosidase assay kit (Mammalian beta Galactosidase Assay Kit, ThermoScientific™) following the manufacturer’s recommendations.

### Lipofuscin Analysis

Lipofuscin analysis was performed with the Sudan Black (SB) dye according to the protocol adapted from Evangelou and Gorgoulis (2017). After preparing the SB solution, the slides were prepared by immersion in decreasing concentrations of xylene and ethyl alcohol and, subsequently, stained with the SB dye. After drying, the slides were mounted with a coverslip in 40% TBS glycerol. Images of the ovarian sections were captured by a digital camera (Moticam 5.0, Motic®, Hong Kong, China) coupled to a microscope (Nikon Eclipse E200, Nikon Corporation, Japan), using 4 and 10X objectives and quantified with the Image J Software.

### Collagen analysis

Collagen analysis was performed using the Picrosirius Red (PSR) dye, which is specific for collagen fibers type I and III (Briley et al., 2016). After preparing the PSR solution, slides were deparaffinized by immersion in decreasing concentrations of xylene and ethyl alcohol and stained with the PSR dye. For PSR staining, slides were immersed in a PSR staining solution prepared by dissolving Sirius Red F3BA (Direct Red 80, CI 357.82, Sigma-Aldrich, St. Louis, MO) in a saturated aqueous solution of picric acid (Sigma-Aldrich, St. Louis, MO) at 0.1% w/v. All slides were processed simultaneously to minimize variation in staining intensity by the automated sample preparation system (ST5020, Leica Biosystems). After drying, the slides were mounted with a coverslip in 40% TBS glycerol. Images of the ovarian sections were captured by a digital camera (Moticam 5.0 Motic®, Hong Kong, China) coupled to a microscope (Nikon Eclipse E200, Nikon Corporation, Japan), using a 10X objective and quantified with the Image J Software.

### Immunofluorescence

Ovarian sections were deparaffinized with xylene and rehydrated with graded alcohols. The anti-CD68 monoclonal antibody (ab955, Abcam, Cambridge, United Kingdom) was diluted in 1.5% BSA solution (Skaznik-Wikiel et al., 2016) at 1:400 dilution. Anti-Lamin B1 (ab133741, Abcam) was also diluted in 1.5% BSA solution and used at 1:1000 dilution (Guo et al., 2014). Antigen retrieval was performed for 3 min at boiling point in citrate solution (pH 6.0). Nonspecific background staining was reduced by covering tissue sections with 10% BSA and 7% goat serum. The slides were incubated overnight with the primary antibody in a humid chamber at 4°C, for 1 h with secondary antibody Alexa Fluor® 488 (ab150077, Abcam) and for 3 min with DAPI (Invitrogen, Carlsbad, USA). Slides were mounted with a drop of mounting medium (Fluoroshield, Sigma-Aldrich, St Lois, USA) under coverslips. For analysis of CD68 and LB1, an area of each section (1 section/mouse) was acquired at 200x magnification using a confocal laser scanning microscope (Leica TCS SP8). Sequential scanning was employed using the 488 nm laser to visualize Alexa 488 and the 405 nm UV lasers to visualize DAPI staining. Fluorescence intensity was calculated using Image J® software to calculate the proportion of positive staining in each ovarian section.

### Statistical Analysis

Statistical analyzes were performed using Graphpad Prism 8 software. Values were tested for normality using the Shapiro-Wilk test. Data was transformed for normality when necessary. Repeated measures ANOVA was used for body weight and ITT analysis. Two-way ANOVA was used for analysis of other continuous variables (follicle count, biochemical/immunofluorescence and oxidative stress). Values of P<0.05 were considered as statistically significant.

## RESULTS

### Body mass and fat content

Females in estropause exhibited higher body mass (Figure 1a) and greater body mass gain post-estropause (Figure 1b) compared to cyclic females. Despite the increase in body mass, there was no difference in visceral fat content between estropause and cyclic mice (Figures 1c). Senolytic treatment did not affect the body mass gain or visceral fat content of either estropause or cyclic mice.

**Fig. 1–.**
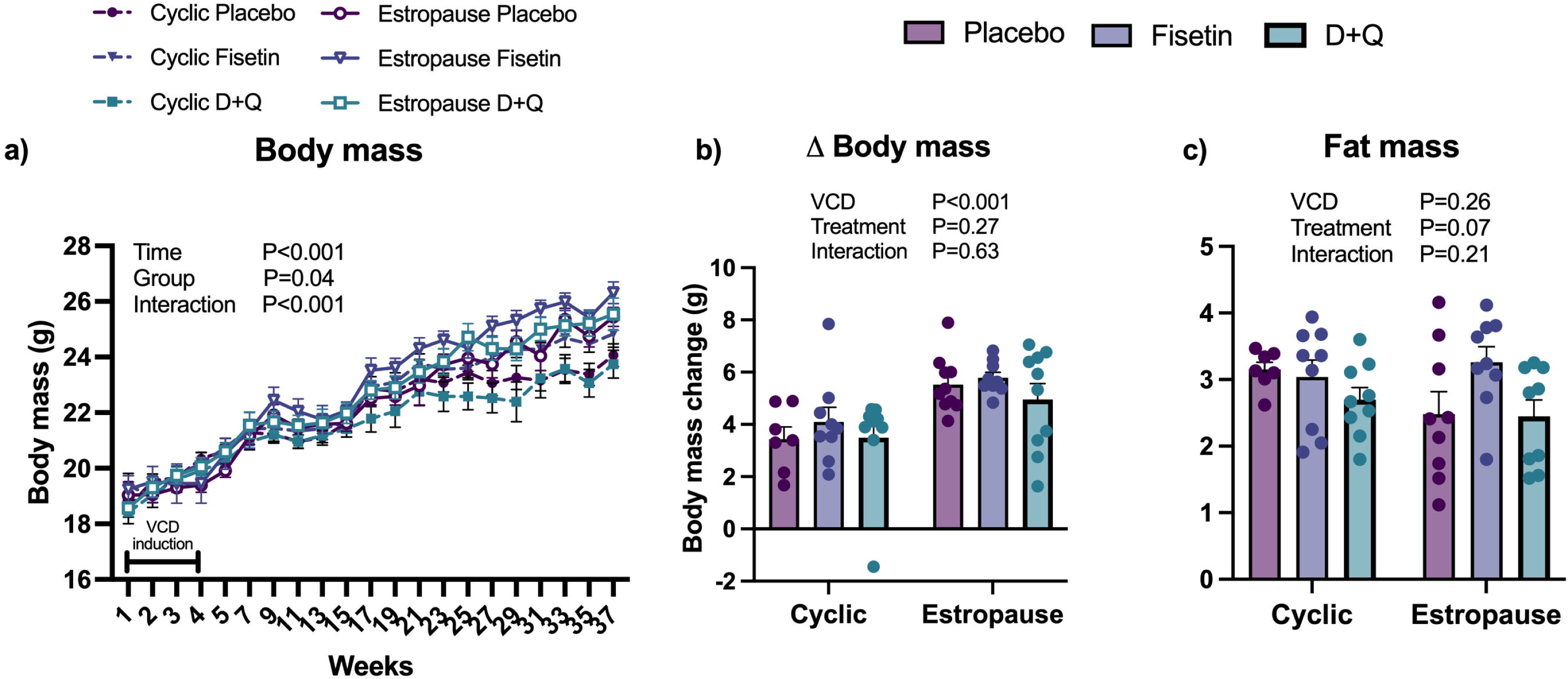
a) Body mass of females from the start of VCD injection to the end of the experimental period; b) body weight gain during the period of senolytic treatment from 5 to 11 months of age; c) Abdominal adipose tissue relative to body mass content.

### Metabolic parameters

There was no difference in insulin tolerance (Figure 2a), basal glucose levels (Figure 2b), and the rate of glucose decay in response to insulin (Figure 2c) between cyclic and estropause females. Senolytic drugs did not affect any parameters of glucose metabolism also.

**Fig. 2–.**
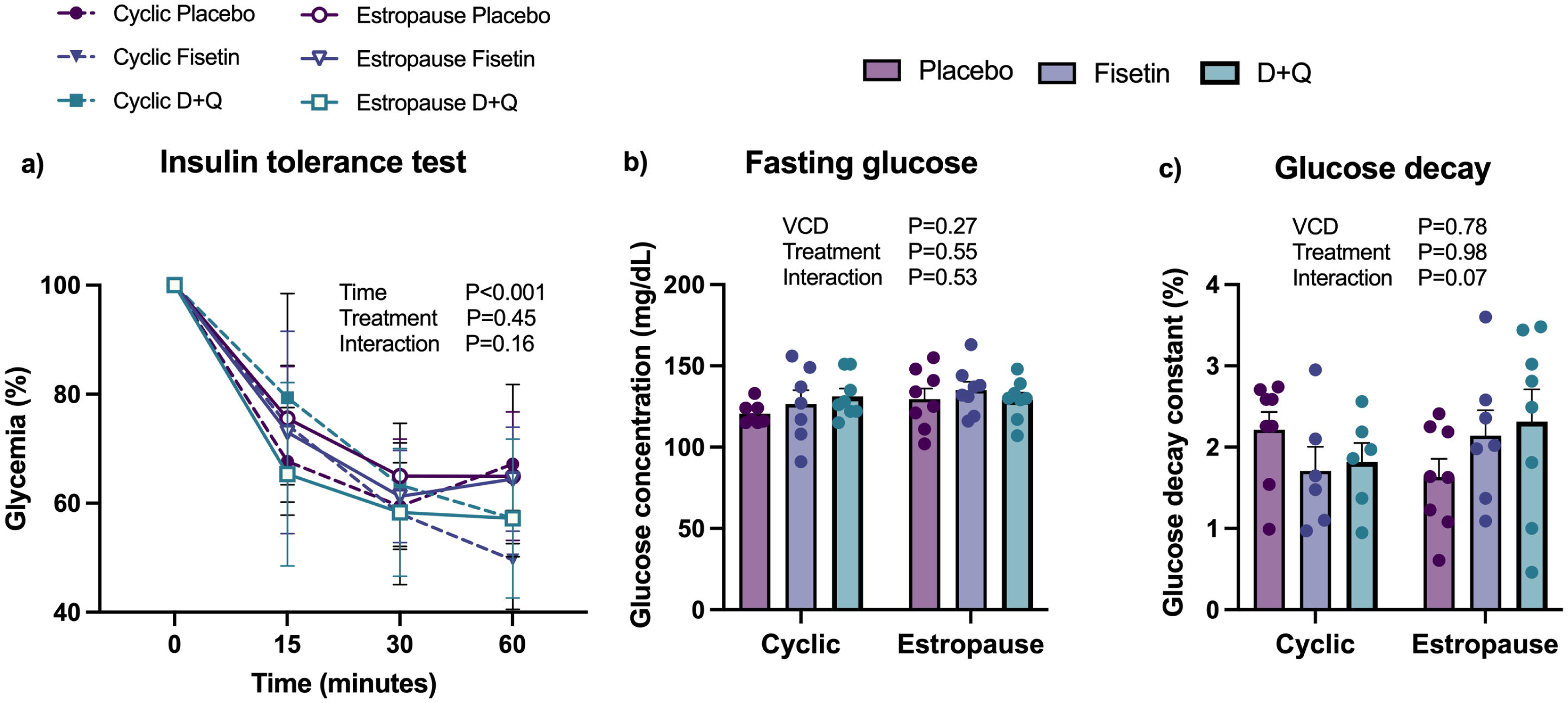
a) Glucose levels during the insulin tolerance test (ITT); b) basal blood glucose levels; c) glucose decay constant during the first 15 min of the ITT.

No differences were found in serum levels of AST and ALT, TP, TC, TG, HDL between estropause and cyclic females. ALB levels were lower in estropause compared to cyclic females (P=0.01). Senolytic drugs did not affect any of the serum biochemical parameters.

### Redox status and cellular senescence in adipose and hepatic tissue

Estropause females exhibited diminished CAT activity in visceral adipose tissue compared to cyclic females (Figure 3d). There was no difference in adipose tissue ROS (Figure 3a), sulfhydryl (Figure 3b), nitrites (Figure 3c) and SOD activity (Figure 3e). In the hepatic tissue, estropause fisetin-treated females had lower ROS levels than placebo estropause females (Figure 4a). VCD tended (P=0.06) to increase SOD activity in liver. No differences were found in the hepatic levels of sulfhydryl (Figure 4b), nitrites (Figure 4c), or CAT activity (Figure 4e). There was no difference in SA-β-gal activity in visceral adipose tissue (Figure 3f) and liver (Figure 4f) between cyclicity status or in response to senolytic drugs.

**Fig. 3–.**
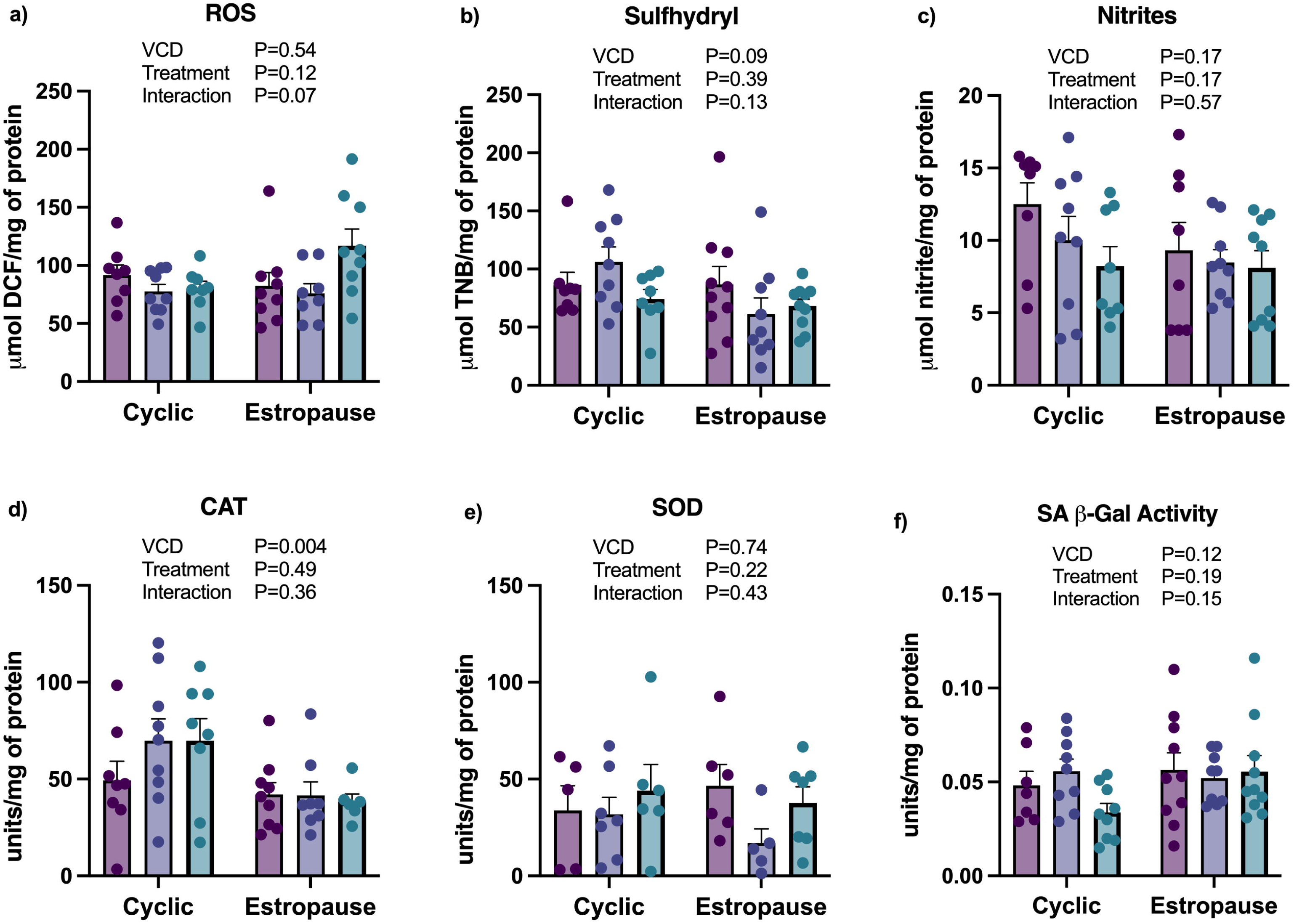
Redox status and senescence in adipose tissue. a) Reactive oxygen species (ROS); b) sulfhydryl (SH); c) nitrites; d) catalase (CAT) activity; e) superoxide dismutase (SOD) activity and f) senescence-associated β-galactosidase (SA-β-gal) activity.

**Fig. 4–.**
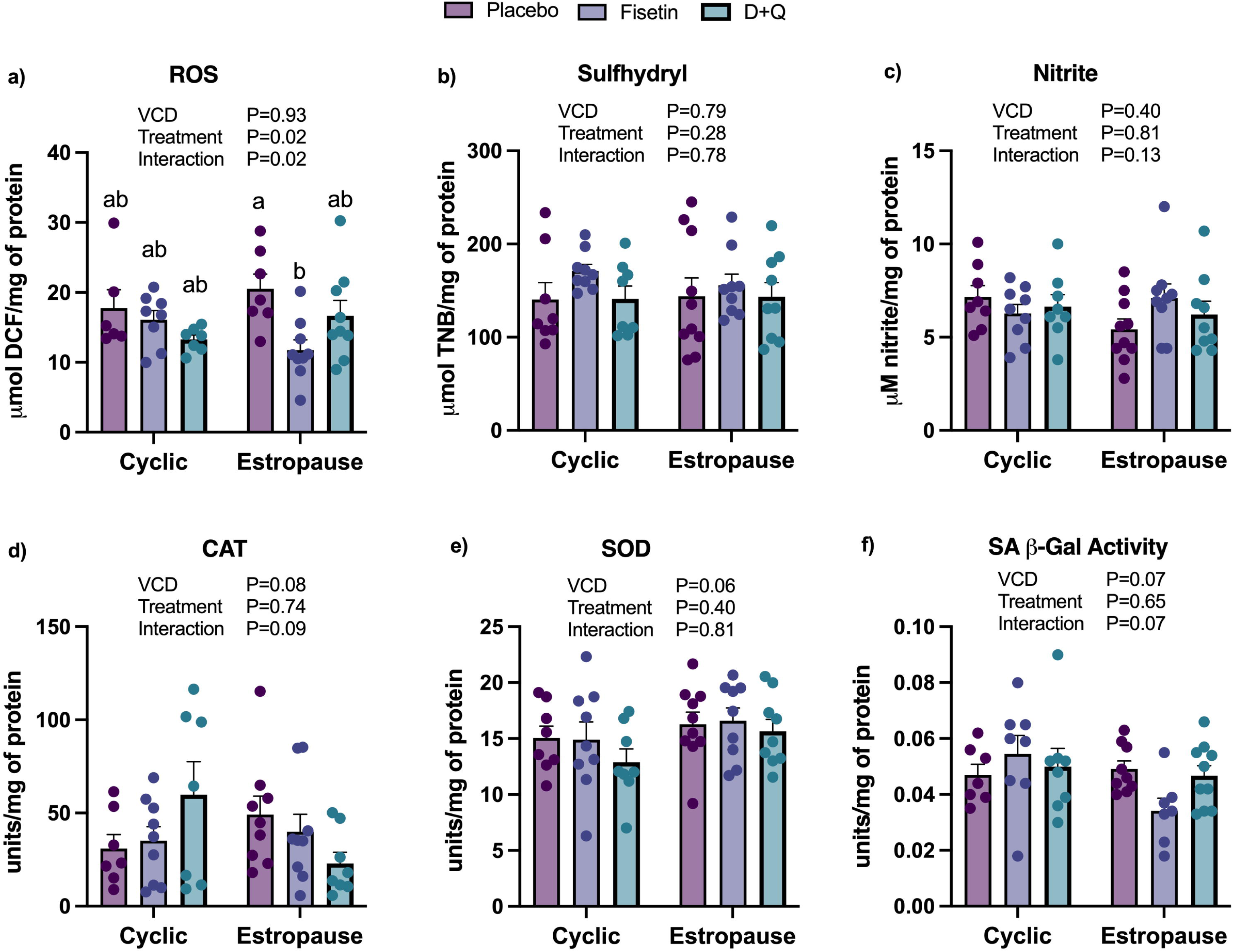
Redox status and senescence in hepatic tissue. a) Reactive oxygen species (ROS); b) sulfhydryl (SH); c) nitrites; d) catalase (CAT) activity; e) superoxide dismutase (SOD) activity and f) senescence-associated β-galactosidase (SA-β-gal) activity.

### Ovarian activity and senescence

The number of ovarian follicles was reduced at all stages of development in estropause compared to cyclic females (Figure 5 a-f). All females injected with VCD were acyclic before starting treatment with senolytics, as confirmed by vaginal cytology. Cyclic females, regardless of senolytic drugs, had similar number of follicles in all stages of development (Figure 5 a-f).

**Fig. 5–.**
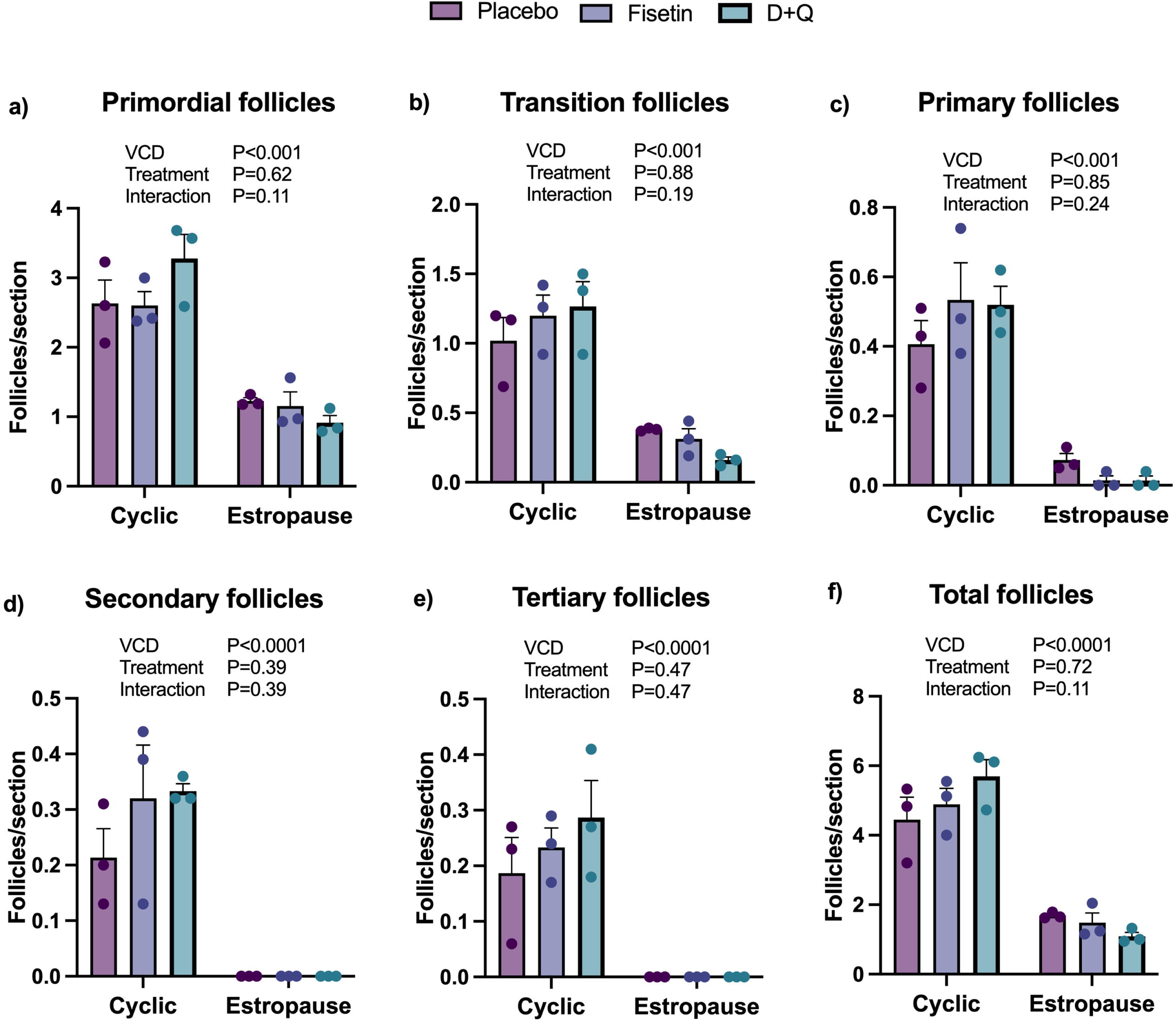
Number of a) primordial, b) transition, c) primary, d) secondary, e) tertiary, and f) total follicles per ovarian section.

Lipofuscin staining was higher in the ovary of estropausal females compared to cyclic females, with no effect of senolytic drugs (Figure 6a). The percentage of lamin-B1 positive cells was reduced in estropausal ovaries compared to cyclic ovaries (Figure 6b), also with no effect from senolytic drugs. Higher lipofuscin and lower lamin-B1 are indicators of increased presence of senescent cells in the ovaries of estropause females. No difference was observed for macrophage infiltration (Figure 6c). We observed that estropause females had increased ovarian fibrosis (Figure 6d); however also without any effects of senolytic drug treatment.

**Fig. 6–.**
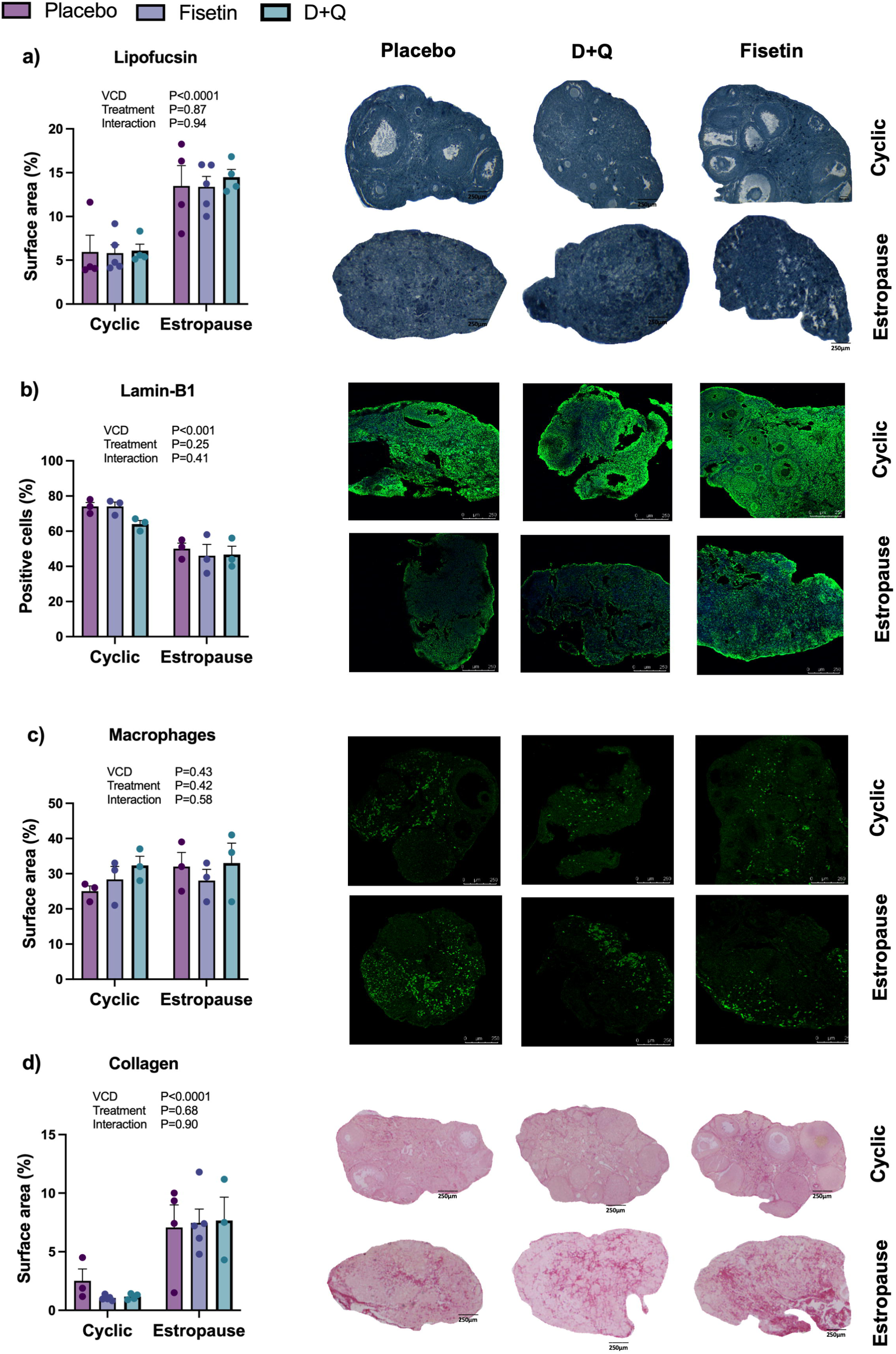
Markers of cellular senescence in the ovary a) percentage of lipofuscin positive cells; b) percentage of lamin B1 positive cells; c) percentage of CD68 poistive cells; d) percentage of PSR staining. Ovarian images representative of each group as follows: 1) cyclic placebo, 2) cyclic fisetin, 3) cyclic D+Q, 4) estropause placebo, 5) estropause fisetin, 6) estropause D+Q.

## DISCUSSION

Treatment with senolytic drugs has been investigated for its potential to selectively clear senescent cells in different tissues, improving health and lifespan (Musi et al., 2018; Kathoon et al.,2021; Bussian et al., 2018; Fang et al., 2023, Luis et al., 2022). However, the effectiveness of these drugs when used in young mice is controversial and seems to be sex specific (Fang et al., 2023; Gao et al., 2023). Aging in females leads to depletion of the ovarian reserve and increased incidence of age-related chronic diseases. Therefore, our aim was to investigate whether inducing estropause in young females could trigger substantial metabolic changes that could be prevented by senolytic therapy. In our study, estropause resulted in increased body mass and only minor metabolic changes. Treatment with fisetin or D+Q did not affect body mass and most metabolic responses in females induced to estropause compared to cyclic females. Estropause females displayed heightened senescence markers in ovarian tissue, which were not affected by senolytic drugs. This suggests that senescent cell burden is confined to the ovarian tissue of young estropause females and does not lead to significant changes in metabolic parameters beyond increased body mass.

The increased body mass in estropausal compared to cyclic females corroborates with our previous study using mice of similar age (Ávila et al, 2023). However, others only observed significant body mass gain in estropause females fed a high-fat diet (Romero–Aleshire et al., 2009), suggesting that estropause alone in mice does not induce many metabolic changes. Senolytic treatment did not affect body mass gain of estropause females. A previous study that evaluated the senolytic potential of D+Q and fisetin in cyclic young females and males only observed beneficial effects in males (Fang et al., 2023). In females, treatment with D+Q induced an increase in body mass gain and fat deposition (Fang et al., 2023), and no effects were observed in fisetin-treated females (Fang et al., 2023). We did not observe increased body mass gain or visceral adiposity in females receiving D+Q, despite females being of similar age to previous studies (Fang et al., 2023). This suggests that the use of senolytic drugs in young females needs to be better understood, as in addition to age, factors such as cyclicity, seems to affect the response to these drugs.

Regarding metabolic parameters, no significant differences were observed between estropause and cyclic females for insulin sensitivity, glycemia, lipid profile, liver injury markers, or TP. Similarly, previous data indicate lower insulin sensitivity only in estropause females exposed to a high-fat diet associated to higher adiposity (Romero-Aleshire et al., 2009; Ávila et al., 2023). This further suggests external influences, such as the type of diet, may have a main role on metabolic changes induced by estropause, and that hormonal changes alone have a small effect. Increased cholesterol level in estropausal females was previously observed, even when fed standard chow (Romero-Aleshire et al., 2009), which was not observed in our study. ALB levels were lower in estropause females compared to cyclic controls. This protein reduces with age in humans (Weaving et al., 2016) and mice (Ding and Kopchick, 2011). Lower ALB levels are related to increased inflammation and cardiovascular disease risk (Gillum, 2000). Therefore, the reduction of ALB levels in estropause females can be an indicator of age-related changes in liver function. Further investigations into metabolic and biochemical alterations in estropause females are warranted to understand how cyclicity status may impact it across different diets. In addition to minimal changes in response to estropause, senolytic treatment also did not influence most of the biochemical parameters analyzed. Changes observed in ALB serum levels in estropausal females were not reversed by senolytic drugs. Therefore, as observed for body mass and adiposity, senolytics exert little influence in the biochemical parameters studied, regardless of cyclicity.

In our study estropause resulted in decreased in CAT activity in adipose tissue, which is noteworthy, given the pivotal role of CAT in the antioxidant defense system (Barbosa et al., 2010). This reduction in CAT activity has the potential to disturb the redox state of adipose tissue. Similarly, a study with ovariectomized rats found reduction in CAT activity in adipose tissue compared to intact controls (Lissarassa et al., 2020). Despite this, other redox parameters in adipose tissue do not indicate a difference between cyclic and estropausal females. In the liver, we observed that ROS levels were higher in liver tissue of estropausal females and SOD activity tended to decrease. SOD1 and SOD2 protein levels do not change in response to ovarian failure in older mice (King et al., 2024). Additionally, senolytics did not affect most of redox state parameters evaluated in our study. Fisetin decreased hepatic ROS levels in estropausal females in comparison to estropausal placebo females, suggesting a benefit from this intervention. Previous findings suggest that D+Q can have some detrimental effects, while fisetin had only limited beneficial effects in young female mice (Fang et al., 2023). Our study is line with these findings and suggest potential benefit for fisetin in some redox parameters. Future studies should focus on longer interventions or much older mice, when redox parameters maybe more compromised.

A previous study showed that senescent cell markers were not affected by fisetin, and some were even increased by D+Q treatment in young females (Fang et al., 2023). We also observed no difference in SA-β-gal activity in adipose and hepatic tissue when comparing cyclic and estropausal females. Senolytic drugs also did not decrease SA-β-gal activity in liver or adipose tissue. This is line with the small benefits observed in redox status of these tissues. SA-β-gal activity is one of the most widely used markers to assess senescent cells both in vitro and in vivo (Dimri et al., 1995; Debacq-Chainiaux et al., 2009; Wang et al., 2018). The lack of effect of senolytic drugs further indicates that young females have little benefits from this therapy. Our observations also suggest that cyclicity independent of age cannot significantly increase senescent cell burden and oxidative stress in liver and adipose tissue. However, further studies evaluating cyclicity status in older females is still warranted.

Next, we aimed to better understand the changes induced by estropause in ovaries, given the significant reduction in the number of follicles resulting from chemical estropause induction. Therefore, we also investigated markers of ovarian senescence, inflammation and fibrosis. Increased lipofuscin (Urzua et al., 2018; Hense et al., 2022) and decreased LB1 (Guo et al., 2014) staining was observed in the ovaries of estropause females in our study. This suggests that estropause induction and the associated follicular depletion increase senescent cell markers in the ovarian tissue. However, senolytic drugs were not able to reduce these senescence markers independent of cyclicity status. Loss of lamin B1 occurs due to chronological aging and accumulation of senescent cells (Wang et al., 2018) or when cells undergo induced senescence in different tissues (Wang et al., 2017; Shimi et al., 2011; Dreesen et al., 2013). Others have shown that young female mice treated with doxorubicin or cisplatin have a severely decreased ovarian reserve and increased markers of cellular senescence (Du et al., 2022; Gao et al., 2023). Senolytics were able to reduce senescence markers in mice receiving doxorubicin and cisplatin (Du et al., 2022; Gao et al., 2023). However, primordial follicles loss was only prevented when senolytics were administered concomitantly to cisplatin (Du et al., 2022). In our study senolytics were only started long after estroupause was confirmed, which could explain why no beneficial effects were observed in senescence markers. This indicates that other strategies may focus on the senolytic therapy to be started before the transition to the estropause state. We also observed increased ovarian fibrosis in estropause compared to cyclic females, which was not reduced by senolytic therapy. Previous studies also observed increased ovarian fibrosis in females induced to estropause with VCD (Sen Halicioglu et al., 2021). Similarly, injection of young mice with doxorubicin or cisplatin promoted ovarian failure and increased ovarian fibrosis (Du et al., 2022; Gao et al., 2023). However, senolytics only prevented ovarian fibrosis when administered concomitantly with cisplatin (Du et al, 2022), similarly to the observed for the ovarian reserve. This suggests that the fibrosis associated with ovarian failure cannot be reversed by senolytic therapy. Others also observed significant more ovarian fibrosis in aged compared to young mice (Landry et al., 2022; Briley et al., 2016), indicating that fibrosis is a product of both age and ovarian failure in young mice. No difference was observed between cyclic and estropause females regardless of treatment for macrophage infiltration (CD68). Others also did not observe a significant increase in ovarian macrophage in females up to nine months of age (Isola et al., 2023).

Although certain studies propose that estropause may be closely linked with metabolic alterations due to the decrease in estrogen production by creating a systemic pro-oxidative state, our findings only indicated an increase in body mass gain. This increase, however, was not accompanied by changes in visceral fat mass nor significant metabolic or oxidative alterations in the liver and adipose tissue. Additionally, we did not observe clear beneficial effects of the senolytics D+Q or fisetin in cyclic or estropause females. There is a need for additional investigation into the effects of senolytics in estropause females. Overall, our current data suggest that estropause was associated to small metabolic changes in young females and that senolytics had no protective effects. Changes in senescence markers were confined to the ovarian tissue of females induced into estropause and remained unaffected by senolytic therapy.

**Table 1–.** Serum biochemical parameters of cyclic and estropause females treated with senolytics dasatinib and quercetin (D+Q) or fisetin (FIS).

| Parameter | Cyclic |  |  |  |  |  | Estropause |  |  |  |  |  | P value |  |  |
| --- | --- | --- | --- | --- | --- | --- | --- | --- | --- | --- | --- | --- | --- | --- | --- |
|  | PLA |  | FIS |  | D+Q |  | PLA |  | FIS |  | D+Q |  | Treatment | VCD | Interaction |
|  | Mean | SEM | Mean | SEM | Mean | SEM | Mean | SEM | Mean | SEM | Mean | SEM |  |  |  |
| **TC (mg/dL)** | 113.2 | 10.4 | 127.7 | 14.7 | 127.8 | 5.8 | 124.8 | 9.7 | 106.1 | 12.4 | 128.8 | 8.5 | 0.52 | 0.73 | 0.30 |
| **HDL**<br>**(mg/dL)** | 40.1 | 2.0 | 48.2 | 3.6 | 42.5 | 1.7 | 43.8 | 3.8 | 40.0 | 3.9 | 41.7 | 3.0 | 0.77 | 0.51 | 0.22 |
| **LDL**<br>**(mg/dL)** | 62.9 | 11.1 | 59.5 | 14.0 | 67.5 | 6.1 | 62.4 | 9.9 | 50.0 | 12.5 | 69.8 | 6.3 | 0.40 | 0.76 | 0.83 |
| **TG (mg/dL)** | 67.4 | 14.2 | 75.6 | 9.2 | 89 | 10.4 | 80.2 | 7.2 | 80.3 | 9.6 | 86.3 | 4.6 | 0.30 | 0.51 | 0.70 |
| **Albumin**<br>**(g/dL)** | 2.5 | 0.07 | 2.4 | 0.1 | 2.5 | 0.1 | 2.2 | 0.1 | 2.3 | 0.1 | 2.3 | 0.1 | 0.97 | 0.01 | 0.81 |
| **Total**<br>**protein** | 5.8 | 0.2 | 5.5 | 0.2 | 5.8 | 0.2 | 5.3 | 0.2 | 5.3 | 0.2 | 5.7 | 0.2 | 0.22 | 0.08 | 0.63 |

| (g/dL) |  |  |  |  |  |  |  |  |  |  |  |  |  |  |  |
| --- | --- | --- | --- | --- | --- | --- | --- | --- | --- | --- | --- | --- | --- | --- | --- |
| **ALT (U/L)** | 43.0 | 4.1 | 38.2 | 2.2 | 37.4 | 1.9 | 39.1 | 2.4 | 39.8 | 2.9 | 40.9 | 2.6 | 0.71 | 0.86 | 0.38 |
| **AST (U/L)** | 254.3 | 24.5 | 211.4 | 13.4 | 183.9 | 22.6 | 211.1 | 18.3 | 217.7 | 21.9 | 227.3 | 14.6 | 0.37 | 0.89 | 0.09 |

PLA (placebo); FIS (fisetin); D+Q (dasatinib +quercetin); VCD (4-vinylcyclohexene diepoxide); TC (total cholesterol); HDL (high-density lipoprotein cholesterol); LDL (low-density lipoprotein cholesterol); VLDL (very low-density lipoprotein cholesterol); TG (triglycerides); ALT (alanine aminotransferase); AST (aspartate aminotransferase).

## ACKNOWLEDGEMENTS

This work was supported by CAPES, CNPq, FAPERGS. This project has been made possible in part by grant number 1023 from the Global Consortium for Reproductive Longevity & Equality (GCRLE).

## AUTHOR CONTRIBUTION

**Bianca M Ávila,** Conceptualization; Data curation; Formal analysis, Writing - original draft **Bianka M Zanini,** Investigation, Visualization, Writing - review & editing **Karina P Luduvico,** Investigation, Visualization, Writing - review & editing **Thais L Oliveira,** Investigation, Visualization, Writing - review & editing **Jéssica D Hense,** Investigation, Visualization, Writing - review & editing **Driele N Garcia,** Investigation, Visualization, Writing - review & editing **Juliane Prosczek, Francielle M Stefanello,** Investigation, Visualization, Writing - review & editing **Pedro Henrique da Cruz,** Investigation, Visualization, Writing - review & editing **Janice L Giongo,** Investigation, Visualization, Writing - review & editing **Rodrigo A Vaucher,** Investigation, Visualization, Writing - review & editing **Jeffrey B Mason,** Conceptualization, Writing - review & editing **Michal M Masternak,** Conceptualization, Funding Acquisition, Writing - review & editing **Augusto Schneider** Conceptualization, Project administration, Funding Acquisition, Investigation, Visualization, Writing - review & editing

